# Chlorotoxin Conjugated with Saporin Reduces Viability of ML-1 Thyroid Cancer Cells In Vitro

**DOI:** 10.1101/2019.12.20.885483

**Authors:** Husref Rizvanovic, A Daniel Pinheiro, Kyoungtae Kim, Johnson Thomas

## Abstract

**Background:** Although differentiated thyroid cancer has good prognosis, radioactive iodine (RAI) resistant thyroid cancer is difficult to treat. Current therapies for progressive RAI resistant thyroid cancer are not very effective. There is an unmet need for better therapeutic agents in this scenario. Studies have shown that aggressive thyroid cancers express matrix metalloproteinase −2 (MMP-2). Chlorotoxin is a selective MMP-2 agonist. Given that Saporin is a well-known ribosome-inactivating protein used for anti-cancer treatment, we hypothesized that Chlorotoxin-conjugated Saporin (CTX-SAP) would inhibit the growth of aggressive thyroid cancer cell lines expressing MMP-2.

**Methods:** The ML-1 thyroid cancer cell line was used for this study because it is known to express MMP-2. ML-1 cells were treated with a toxin consisting of biotinylated Chlorotoxin bonded with a secondary conjugate of Streptavidin-ZAP containing Saporin (CTX-SAP) from 0 to 600 nM for 72 hours. Then, cell viability was measured via XTT assay at an absorbance of A_**450-630**_. Control experiments were set up using Chlorotoxin and Saporin individually at the same varying concentrations.

**Results:** After 7 hours of incubation, there was a statistically significant reduction in cell viability with increasing concentrations of the CTX-SAP conjugate (F=4.286, p=0.0057). In particular, the cell viability of ML-1 cells was decreased by 49.77% with the treatment of 600 nM of CTX-SAP (F=44.24), and the reduction in cell viability was statistically significant (Dunnett’s test p<0.0001). In contrast, individual Chlorotoxin or Saporin in increasing concentrations had no significant effect on cell viability using similar assay.

**Conclusion:** This *in vitro* study demonstrated the efficacy of a CTX-SAP conjugate in reducing the viability of ML-1 thyroid cancer cells in a dose dependent manner. Further studies are needed to delineate the effectiveness of CTX-SAP in the treatment of aggressive thyroid cancer. Our study points towards MMP-2 as a potential target for RAI-resistant thyroid cancer.

## Introduction

Incidence of thyroid cancer has been increasing in the past decade. Estimated incidence of thyroid cancer in 2019 is 52,070 (1). It is predicted that by 2030, thyroid cancer will be the fourth leading cause of new cancer diagnosis in the United States (2). In 2016 there were 822,242 patients living with thyroid cancer in the United States (1). Most of the thyroid cancers respond well to surgery, radioactive iodine, and thyroid stimulating hormone (TSH) suppression. However, a subset of these thyroid cancers will develop metastasis and become radioactive iodine (RAI) resistant. According to a study by Schlumberger *et al*., up to 50% of thyroid cancer with metastasis may develop inability to concentrate iodine (3). When thyroid cancer cells become resistant to RAI, newer therapeutic agents like tyrosine kinase inhibitors could be used. Even with these newer therapeutic agents, most RAI resistant metastatic thyroid cancer will progress. This could also result in multiple toxicities. Even with newer therapeutic agents average progression free interval is about 18 months (4) and the ten year mortality rate for these kinds of thyroid cancers can reach up to 50% (5). Hence, there is an unmet need for better therapeutic agents for RAI resistant thyroid cancer. In this paper, we explore the effect of Chlorotoxin-Saporin conjugate on the ML-1 thyroid cancer cell line.

Chlorotoxin was initially identified in the venom of scorpion *Leiurus quinquestriatus* (6). It binds specifically to isoform 2 of matrix metalloproteinase (MMP-2) (7). MMP-2 is overexpressed in aggressive thyroid cancers when compared to normal thyroid tissue (8). Tumors with larger size, extrathyroidal invasion and lymph node metastasis were found to overexpress MMP-2(9). Papillary thyroid cancers (PTC) with lymph node metastasis were found to have significantly higher ratio of pro-MMP-2 activation when compared to follicular adenomas and normal thyroid tissue (10). Widely invasive follicular thyroid cancer was also found to express MMP-2 (11).

Saporin (SAP) is a ribosome-inactivating protein (RIP) derived from the seeds of the soapwort plant, *Saponaria officinalis* (12). The mechanism of action of Saporin has been well studied and successfully employed in the creation of immunotoxins. Saporin is very stable *in vivo* and is resistant to proteases in the blood. Since Saporin works through many different cell death pathways, it is hard to develop resistance to it (13–15). Saporin by itself is unable to cause significant cell damage, since it cannot enter the cell efficiently (15). Conjugation with antibodies or other toxins which promotes its internalization, confers lethality to SAP. The first study using an antibody conjugated with SAP was conducted in humans for refractory Hodgkin’s disease, in which 75% of the patients achieved complete remission and 50% of them experienced relief from symptoms (16). A recent study used Substance P conjugated with SAP intrathecally in cancer patients with intractable pain (17).

ML-1 (ACC-464), thyroid cancer cell comes from dedifferentiated recurrent follicular thyroid cancer from a 50 year old patient (18). ML-1 is tumorigenic in rodents. Grimm *et al*., demonstrated that ML-1 cell lines express MMP-2 (19). In this study we assessed the effect of CTX-conjugated with SAP (CTX-SAP) on cell viability of ML-1 cells.

## Materials and methods

### Toxins and Reagents

The Chlorotoxin-Saporin (BETA 010) conjugate was acquired from Advanced Targeting Systems (San Diego, CA). This toxin consisted of biotinylated Chlorotoxin bonded with a secondary conjugate of Streptavidin-ZAP containing Saporin. Unconjugated Saporin and Chlorotoxin were acquired through Sigma-Aldrich. Dulbecco’s Modified Eagle Medium (DMEM), Fetal Bovine Serum (FBS), Phosphate Buffered Saline (PBS), Trypsin-EDTA (and unconjugated Trypsin), Propidium Iodide (PI), and Penicillin-Streptomycin antibodies were ordered from Fischer Scientific. XTT activation reagent (PMS) and solution were purchased from Biotium.

### Thyroid cancer cell line

ML-1 (ACC-464) thyroid cancer cells were acquired from the DSMZ German Leibniz Institute of Microorganisms and Cell Cultures. ML-1 cells were cultured in DMEM supplemented with 10% FBS and 1% Penicillin-Streptomycin antibodies in a 75 cm^3^ Corning culture flask at 37 °C and 5% CO_2_.

### Toxin Treatment

Cells were seeded at a density of 7500 cells/well on a 96-well plate and given 24 hours to incubate and attach to the 96-well plate. Then, varying amounts of CTX, SAP, and CTX-SAP were treated in triplicate or quadruplet repeats to the cells with an increasing dosage, ranging from 0 (NTC) to 600 nM, by using 2 μM stock solutions for CTX, SAP, and CTX-SAP. The final volume of media including the toxin treatment per each well was 100 μL. This was incubated for a period of 72 hours.

### Cell Viability

XTT (2,3-Bis-(2-Methoxy-4-nitro-5-sulfophenyl)-2H-tetrazolium-5-carboxanilide, disodium salt) assay is based off the cleavage of the tetrazolium salt. XTT, in the presence of N-methyl dibenzopyrazine methyl sulfate (PMS), an electron-acceptor, is reduced to form a water soluble, orange-colored formazan salt (20). This type of reaction can only occur in viable, metabolically-active cells, and therefore, the amount of dye formed is directly related to the number of metabolically active cells present (21). In this study, after a 72 hour incubation period with toxin treatment, 25 μL of activated XTT dye was added to each well containing 7,500 cells according to the manufacturer’s guidelines without removing media, and the level of color change was quantified using a multi-well spectrophotometer (A_**450-630**_) (22). This setting was chosen as the Biotium manufacturer suggested the signal absorbance be read at 450-500 nm, and have it be subtracted from the background at 630-690 nm. This is done by setting the plate-reading spectrophotometer to Delta.

### Statistical analysis

Prism8 statistical software (GraphPad Software Inc.) was utilized to conduct statistical analysis. One-Way ANOVA was performed to compare absorbance values between toxin-treated groups and non-treated controls. Post hoc comparisons were done using Dunnett’s test to compare values from toxin-treated groups to the NTC. All values are expressed as the mean and standard deviation of recorded absorbance values. P-values calculated from Dunnett’s test were adjusted to account for multiple comparisons.

## Results

### Unconjugated chlorotoxin (CTX)

ML-1 cells were exposed to unconjugated chlorotoxin at concentrations ranging from 0 to 600 nM for 72 hours (Fig. 1A). There was an overall statistically significant difference in cell viability assessed after 7 hours of incubation in the presence of XTT and PMS (F=3.34, p=0.038). The largest difference was slightly increased viability (absorbance) of the cells exposed to 600 nM of unconjugated CTX relative to the non-treated control (mean difference in absorbance - 0.054). However, this was not statistically significant with a Dunnett’s test (p=0.1279).

**FIG. 1A.**
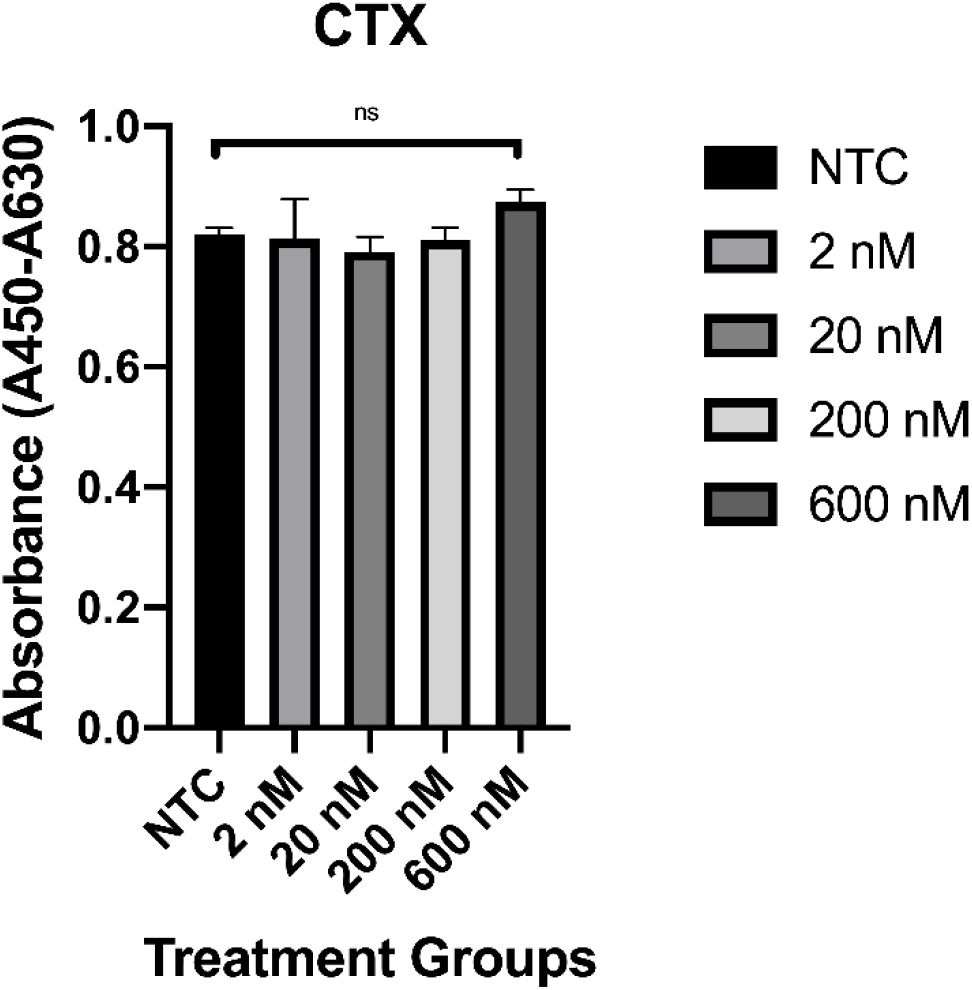
XTT assay results for unconjugated Chlorotoxin. Representation of cell viability with differing amounts of unconjugated Chlorotoxin ranging from 0 nM (NTC) to 600 nM. Error bars represent the standard deviation of each sample. There was no significant effect on ML-1 cell viability.

### Saporin alone (SAP)

ML-1 cells were exposed to unconjugated Saporin at concentrations ranging from 0 to 600 nM (Fig. 1B). There was an overall statistically significant difference in cell viability after 7 hours of incubation with XTT and PMS (F=3.271, p=0.0407). The largest difference was improved viability (absorbance) of the cells exposed to 20 nM of unconjugated SAP relative to the non-treated control (mean difference in absorbance −0.05325). However, this was not statistically significant with a Dunnett’s test (p=0.0874).

**FIG. 1B.**
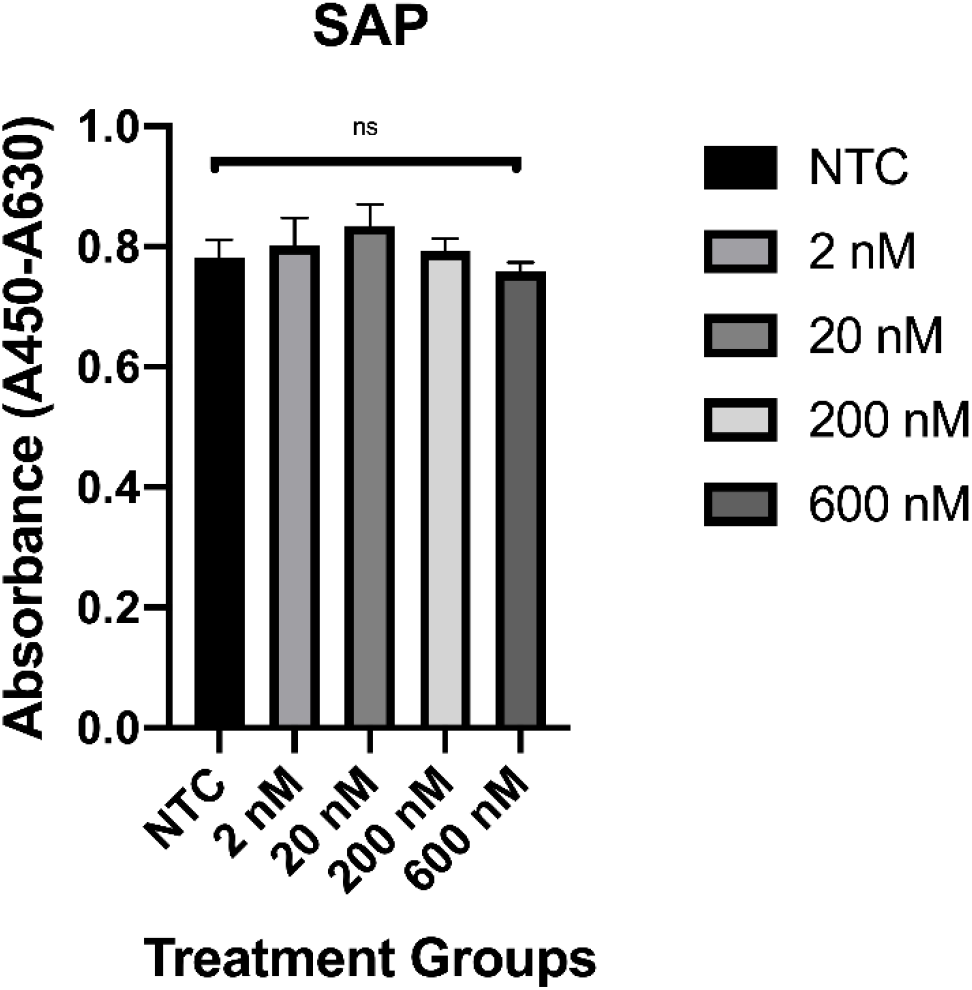
XTT assay results for unconjugated Saporin, at a concentration of 0 nM to 600 nM, on ML-1 cell proliferation. There was no significant effect on cell viability of ML-1 cells.

### Chlorotoxin-Saporin Conjugate (CTX-SAP)

ML-1 cells were exposed to Chlorotoxin-Saporin conjugate at concentrations ranging from 0 to 200 nM (see Fig. 2A). There was an overall statistically significant difference in cell viability at 7 hours of incubation with XTT and PMS (F=4.286, p=0.0057) with an apparent trend for decreased viability with increasing concentration of the conjugate. Post hoc statistical comparisons using Dunnett’s test showed no significantly reduced viability for cells exposed to 2, 10 or 20 nM relative to non-treated control (p>0.05). However, cells exposed to CTX-SAP conjugate at concentrations of 40, 100 and 200 nM had significantly reduced viability relative to non-treated controls (Dunnett’s tests, p=0.0138, p=0.0052, and p=0.0037 for 40, 100 and 200 nM, respectively, relative to NTC).

**FIG. 2A.**
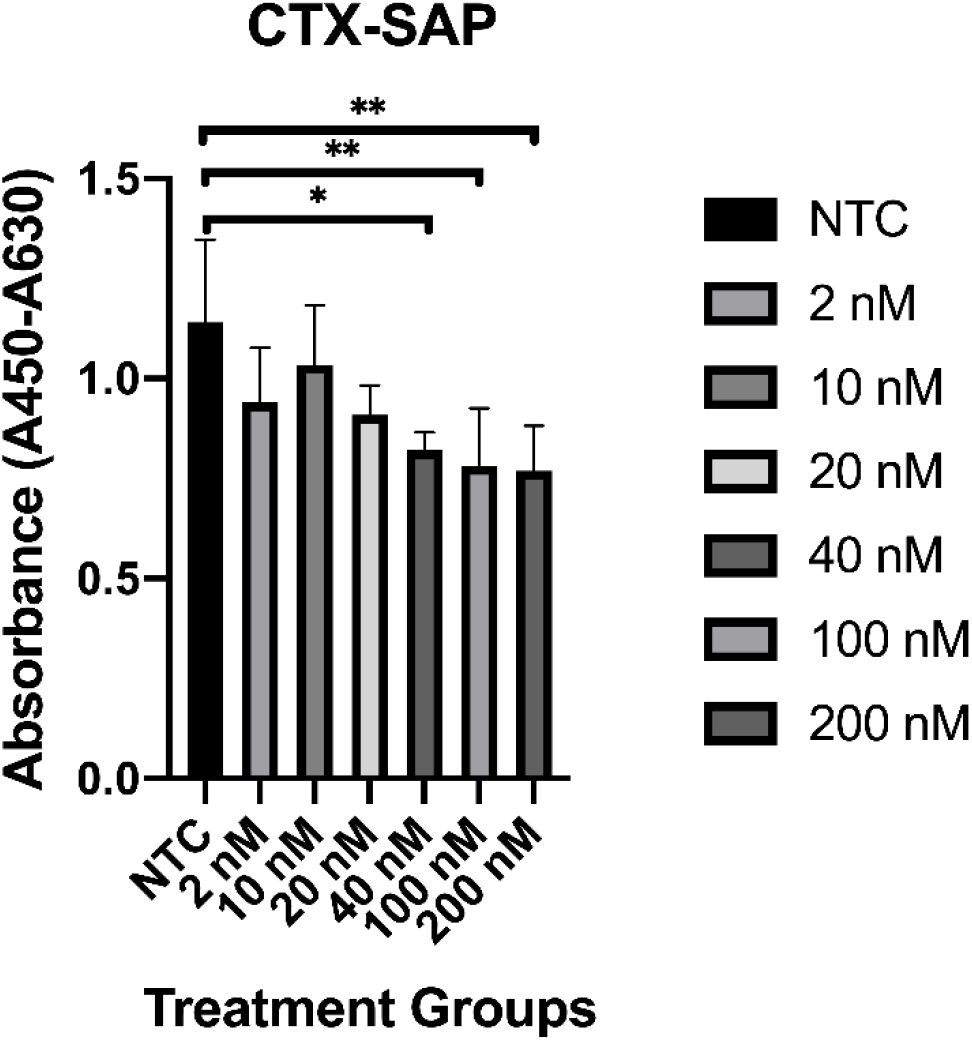
XTT assay results for Chlorotoxin conjugated with Saporin on ML-1 cell proliferation. Dose-dependent inhibition of ML-1 thyroid cancer cell proliferation with the CTX-SAP conjugate treatment at varying concentrations up to 200 nM.

We repeated the experiment with a higher concentration of the CTX-SAP conjugate and once again assessed viability at 7 hours of incubation (Fig. 2B). We used concentrations of 2, 20 and 600 nM, and there was a statistically significant difference in cell viability assessed with a one-way ANOVA (F=44.24, p<0.0001). Post hoc comparisons using Dunnett’s test revealed that viability was significantly reduced for the 600 nM group relative to the non-treated control (difference in absorbance of 0.8410, p<0.0001). However, the lower concentrations of the conjugate did not significantly differ from the non-treated control (p=0.2492 and p=0.5658 for 2 and 20 nM, respectively).

**FIG. 2B.**
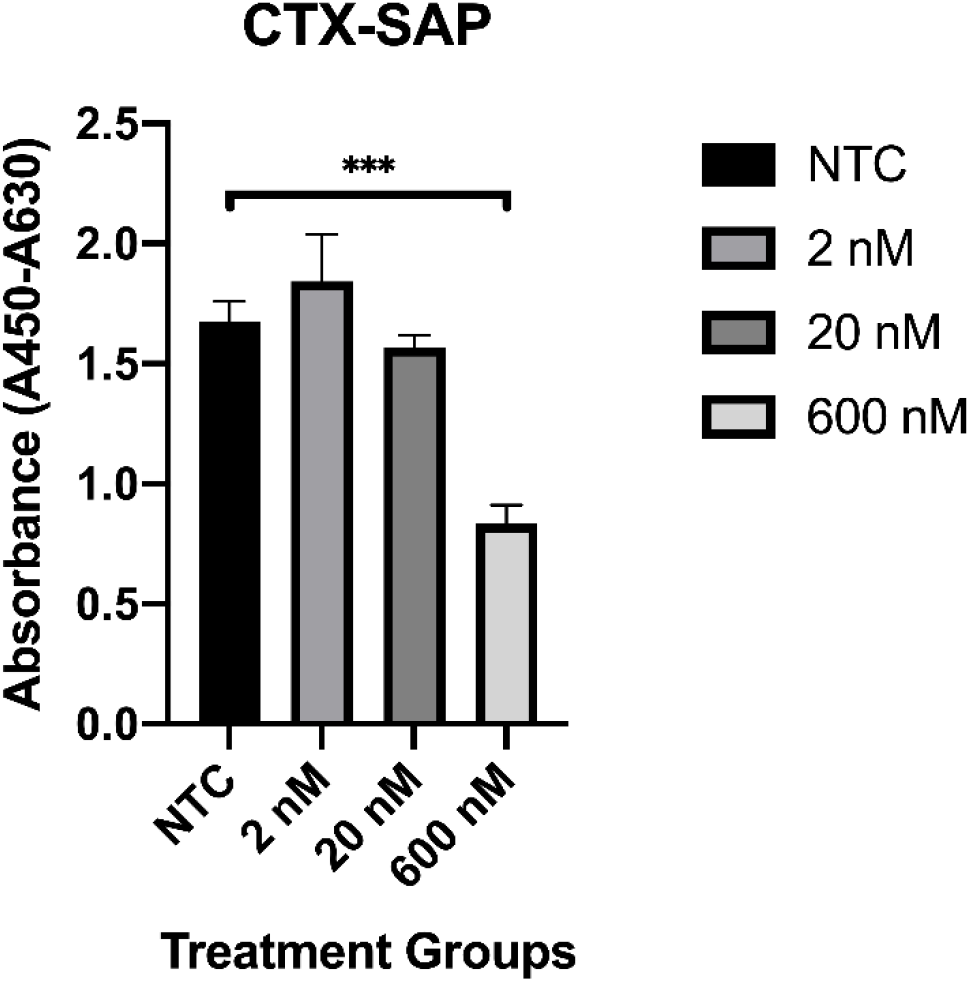
Viability of ML-1 cells was decreased by 49.77% with treatment of 600 nM of Chlorotoxin Saporin conjugate.

## Discussion

This study addressed the effectiveness of Saporin, Chlorotoxin and CTX-SAP conjugate in decreasing cell viability of ML-1 cells. Saporin, a toxin from the plant seed *Saponaria officinalis*, inhibits proliferation or cell viability of cancer cells (23). Previous studies have proven that Saporin is an effective ribosome-inactivating protein (RIP) that inhibits protein synthesis and growth of both normal and tumor cells (24). However, our data shows that there were no significant effects on ML-1 cancer cell viability when treated with unconjugated Saporin up to 600 nM **(Fig. 1A)**. This would be in agreement with other studies that have chosen to use Saporin conjugates for cancer therapies rather than unconjugated Saporin alone (12). This data suggested the need for a vehicle that can be conjugated to Saporin and help the toxin be internalized. Chlorotoxin, which is a 36 amino-acid peptide from the venom of *Leiurus quinquestriatus*, has been previously used as a vehicle to deliver anti-cancer drugs to cancer cells (25). Chlorotoxin is also a known MMP-2 isoform agonist, that makes an effective vehicle for internalization as most aggressive thyroid cancers express MMP-2 receptors (7).

MMP-2 belongs to the matrix metalloproteinases family which helps in degradation of basement membranes and extracellular matrix. MMP-2 is also associated with angiogenesis inside the tumor (26). This promotes spread of the cancer. Activation of MMP-2 can occur at the cell membrane. MMP-2 production is enhanced in papillary thyroid cancer (10, 27, 28). Studies have shown direct correlation between the expression of MMPs and tumor invasion and metastasis (29). Aggressive variants of thyroid cancers tend to express more MMP-2 (10). Thyroid cancer cell lines - BCPAP (poorly differentiated thyroid cancer), K1 (papillary thyroid cancer), CGTH-W-1 (follicular thyroid cancer with metastasis to sternum) and FTC133 (follicular thyroid cancer) also express MMP-2 (19, 30). Anaplastic thyroid cancer cell line SW 579 also express MMP2 (31). Increased expression of MMP-2 is also seen in urothelial, prostate, breast, stomach cancers and in gliomas. Chlorotoxin has been shown to help in the endocytosis of MMP-2(7). Study by Kalhori and Törnquist demonstrated that MMP-2 is involved in ML-1 cell line’s invasive potential (32).

In our study, unconjugated chlorotoxin did not show any significant reduction in cancer cell viability when compared to untreated controls **(Fig. 1B)**. It was hypothesized that a Chlorotoxin and Saporin conjugate would inhibit the cell growth of ML-1 thyroid cancer cells. We observed a significant dose-dependent inhibition of ML-1 cell viability when treated with 40-600 nM of conjugated toxin (**Fig. 2A)**. The most statistically significant reduction of ML-1 thyroid cancer cell viability occurred with treatment of 600 nM of CTX-SAP. Cell viability was decreased by 49.77% with 600 nM of CTX-SAP when compared with non-treated controls **(Fig. 2B)**.

Further studies are needed before this can be used in clinical practice. This study needs to be replicated in other thyroid cancer cell lines also. Reliable methods to detect the presence of MMP-2 in cancer cell lines will help us identify potential malignancies that can be treated with CTX-SAP. Radioiodine[**^131/125^I**] labelled synthetic Chlorotoxin has been successfully administered without any significant side effects *in vivo* (33–36). This could be used to image tumors that are RAI-resistant but express MMP2. Further studies are also needed to understand the effect of this toxin on tumor microenvironment and surrounding normal cells.

## Conclusion

Our study demonstrated that CTX-SAP decreased cell viability of ML-1 cells which express MMP-2. This study points towards MMP-2 as a potential target for RAI-resistant thyroid cancer. Further studies are needed to develop safe and effective treatment against aggressive thyroid cancer using CTX-SAP.

## Acknowledgments

We thank the Department of Biology at Missouri State University, Springfield, Missouri, United States for facilitating this study. Additionally, we are indebted to Hazzar Abysalamah, Hanna Williams, and Dr. Christopher Lupfer for help with cell culture.

